# Deep Recurrent Q-Learning Captures the Behavioral Dynamics Observed in Deterministic and Stochastic Task Switching

**DOI:** 10.64898/2026.03.10.710844

**Authors:** Andrew H. Fagg, Martin Diges, Abigail Zdrale Rajala, Golnaz Habibi, Aaron J. Suminski, Luis Populin

## Abstract

Cognitive Flexibility (CF) is the ability to switch between tasks, even under conditions when the need to switch is not explicitly cued. While the prefrontal cortex and its interaction with subcortical regions are considered central to CF, a key question remains: what are the underlying computational mechanisms that implement the switch from one task to another? In particular, does the switch rely on 1) learning processes in which synaptic changes directly alter action execution choice, or 2) neural state processes that estimate a belief state from which actions can be chosen? Bartolo and Averbeck (2020) argue for the neural state change hypothesis, proposing a Bayesian *belief state* estimation model, and ruling out Reinforcement Learning as an approach to modeling CF tasks because of its reliance on synaptic changes to implement the switch.

We propose instead a Reinforcement Learning-based Deep Recurrent Q-Learning (DRQL) model that simultaneously learns to update a belief state representation based on prior action outcomes, and an action preference representation based on this belief state. This model is presented with a repeated-trial, force-choice probability switching task (PST) in which actions are rewarded stochastically, and the reward probabilites switch between blocks of trials. Although the model is not explicitly cued to the task type, probability of reward, or time of switch, following training, the model performs the PST in the absence of synaptic changes. We show that the trained model produces behavior consistent with non-human primates performing a similar task, and that it develops a belief state representation that captures key information about the current state of the task.

**Significance Statement:** The proposed DRQL model learns to perform the PST, even when reward probabilities vary across different blocks of trials. Behaviorally, the trained model requires different amounts of time to commit to a task switch, depending on the uncertainty of the reward information, with quicker switches for more certain outcomes. The learned belief state and other computed measures may give insight into the neural mechanisms underlying task switching in the primate brain. In contrast with other approaches, the DRQL model presents a more biologically tractable solution to CF, as it is easily adapted to other forms of the PST task, including altering the number of possible actions and the rules for providing rewards.

## Introduction

Cognitive flexibility (CF) is the ability to switch one’s responses to adapt to changing situations, particularly when the need to switch is not explicitly cued (Mansouri et al., 2006; Avila et al., 2015; Cole, 2024). CF is typically studied with reversal learning tasks of varying complexity (Mitz et al., 1991; Dias et al., 1996; Swainson et al., 2000; Cools et al., 2001; Rygula et al., 2010; Rudebeck et al., 2013; Costa et al., 2015; Farashahi et al., 2017), which entail learning associations between stimuli and motor responses, and then adapting appropriately to uncued changes in this association. Typically, subjects receive sparse outcome information for each response (e.g., a reward). The prefrontal cortex (PFC) has long been considered central to CF, as evidenced by PFC lesions that impair task switching (Milner, 1963; Dias et al., 1996), and the observation that PFC neurons encode outcome signals important for updated behavior (Mansouri et al., 2006; Watanabe, 1989; Niki and Watanabe, 1979; Rajala et al., 2020). However, it is increasingly clear that CF depends on broader cortical/subcortical networks that interact dynamically with PFC (Rajala et al., 2020; Birn et al., 2019; Cole, 2024; Costa et al., 2019). The computation implemented through their interaction to support CF tasks remains poorly understood.

Solving deterministic switching tasks in which there are two action choices requires only a memory of the last action and knowledge of its outcome. In stochastic tasks with probabilistic rewards, however, failing to receive a reward could result from either an incorrect or correct action. Thus, identifying the time of switch requires computational machinery that can integrate action/outcome information over time. Bartolo and Averbeck (2020) contrast two computational hypotheses for how switching is implemented in over-trained animals: 1) a Reinforcement Learning (RL)-based model that relies on synaptic changes to implement the switch, and 2) a hand-designed Bayesian-based model that estimates a *belief state* about the current task, and then acts accordingly. The time required to switch in the former model is dominated by synaptic dynamics (i.e., a “learning rate” parameter), whereas this time can vary for the latter model, depending on outcome ambiguity. Because NHPs exhibit switching behavior consistent with the latter, Bartolo and Averbeck (2020) conclude that the RL-based model is insufficient.

Zhang et al. (2025), however, caution about ruling out RL as a hypothesis based on a specific implementation because *RL methods* represent a large class of approaches that vary in their assumptions and implementation details. Here, we argue that the key distinction for RL implementations in CF tasks rests on whether a task switch relies on processes that require: 1) synaptic changes, or 2) neural state changes. While we concur with Bartolo and Averbeck (2020) that the neural state change hypothesis better explains switch timing variability, we propose a RL-based approach that can 1) learn to perform novel tasks and 2) implement task switching based solely on neural state changes. Specifically, Deep Recurrent Q-Learning (DRQL) models couple a recurrent neural network (RNN) to implement belief state estimation with a second neural network (NN) to estimate the value of taking an associated action (Boutilier and Poole, 1996; Rao, 2010; Huang and Rao, 2013; Lambrechts et al., 2022a; Rao, 2024). This approach has the advantage of not relying on hand-designed rules for either constructing the belief state or for choosing actions.

We demonstrate this idea within a probability switching task (PST) in which both NHPs and an artificial agent perform the task under deterministic and stochastic reward conditions. None of the agents is explicitly cued with task type, reward probability, or switch time. We hypothesize that a DRQL-based model that simultaneously learns belief state and action value representations will 1) consistently learn to perform the PST task, 2) produce behavior similar to that of NHPs, and 3) construct a belief state representation that captures the expected reward probability and an estimate of the ideal action choice.

## Materials and Methods

In this paper, we present results from parallel empirical and modeling efforts for a PST. We first describe the NHP PST behavioral paradigm, and then present the structure and training procedure for the DRQL-based model.

### Experimental Design

*Subjects*: Three adult male Rhesus monkeys (Macaca Mulatta) weighing 12 kg (GD), 10 kg (M), and 14 kg (G) were used in this study. The Rhesus monkey was selected because the organization of its brain, as the frontal lobe is the closest homolog to the human among the animal models available for studies of this type (Rudebeck et al., 2019; Wise, 2008; Preuss and Wise, 2022). These nonhuman primates (NHPs) were implanted with a magnetic resonance imaging (MRI) compatible head post to restrain the head for oculomotor and electrophysiological recordings as described earlier (Rajala et al., 2020). All procedures were approved by the University of Wisconsin-Madison Animal Care and Use Committee (IACUC), and were in accordance with the National Institutes of Health *Guide for the Care and Use of Laboratory Animals*.

The NHPs were first trained to earn water rewards for making saccadic eye movements to spots of light presented on a computer screen. Subsequently, they were trained to perform a two alternative forced choice task called probability switching task (PST) that required the selection of one of two targets, a square and a circle, presented simultaneously, 10 deg to the left and right of a central fixation point with a saccadic eye movement (Figure 1). A trial began with the presentation of a fixation red dot at the center of a computer screen, which the subject had to acquire and fixate for 300 − 500 ms. Next, the targets were presented for 800 − 1300 ms. The offset of the fixation dot served as the command for the subjects to select one of the two targets with a saccadic eye movement. For each trial, the left/right presentation of the circle and the square were randomly determined.

**Figure 1:**
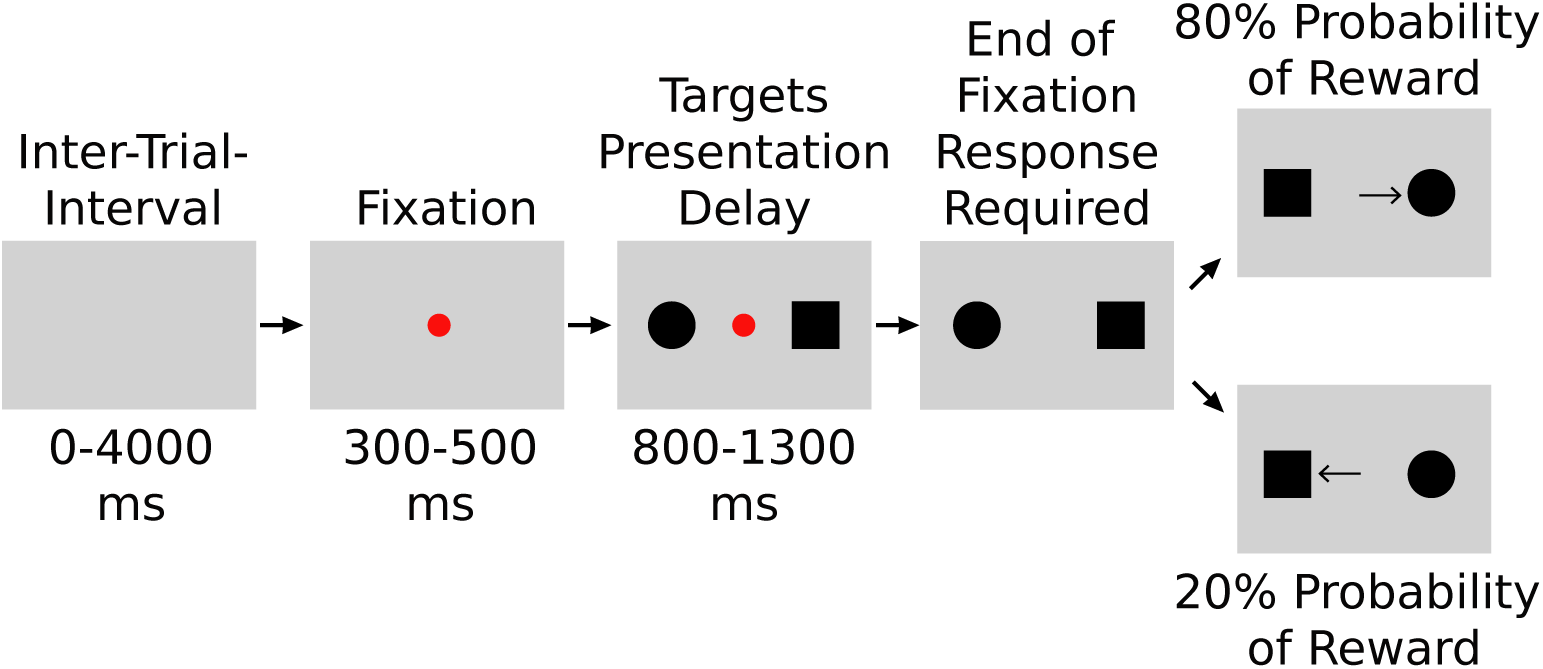
Representation of the Probability Switching Task (PST). Nonhuman primates were presented with a fixation target in the form of a red dot at the center of a computer screen. After a variable period of fixation lasting 300-500 ms, two figures, a circle and a square, were presented 10 deg to the left and right of the fixation point, centered at 0 deg elevation. The left/right location of the targets was determined randomly for each trial. For each block of 100 trials, the two targets were assigned probabilities of reward: 100/0; 90/10; and 80/20. In this example, an 80% probability of receiving a reward was assigned to the circle, and 20% to the square. Upon extinguishing the fixation point, which constituted the signal to respond, the nonhuman primates had to make their selection by making a saccadic eye movement to the chosen target. At the completion of the 100 trials comprising a block, the probability assignments were reverse.

In the example shown in Figure 1, the circle is rewarded with a probability of 80% and the square 20%. At the completion of a block of 100 trials, the probabilities are reversed, and the subjects are expected to adjust their behavior accordingly. The monkeys are tested with 100*/*0; 90*/*10; and 80*/*20 probability schemes. In the 100*/*0 probability scheme (the deterministic configuration), selection of one target always yields a reward, with the other target yielding no reward. In the 90*/*10 and 80*/*20 probability schemes (the stochastic configurations), selection of a target is rewarded with probabilities 90% and 80%, respectively, with the other target yielding rewards of 10% and 20% of trials. The magnitude of the rewards is identical for both high and low probability targets. For any given block of trials, we refer to the selection of the target with the highest probability of reward as a *correct* response.

One subject (GD) was tested in the laboratory, sitting upright in a primate chair facing a computer screen positioned 84 cm in front of the eyes. The second and third subjects were tested during task-based functional Magnetic Resonance

Imaging (fMRI) sessions, placed in the sphinx position in the bore of the magnet. As in Birn et al. (2019), the NHPs were inserted into the bore of the magnet with their heads pointing outward. The task stimuli were presented on a MR-compatible computer screen, positioned 150 cm in front of the subjects’ eyes. A MR-compatible spout placed immediately in front of the subjects’ lips was used to deliver water rewards when appropriate, based on both the subjects’ selections and the probability scheme.

*Eye Movement Recordings and Behavioral Data Analysis*: A video-based eye tracker (Eyelink 1000Plus, SR Research, Ottawa, Canada) was used to record the subjects’ eye movements, during the performance of the PST. Data from a session configured with a 80*/*20 probability scheme collected in the bore of the MRI scanner are shown in Figure 2. The horizontal component of saccadic eye movements was used to select one of the two targets presented to the left and right of the fixation point. This eye position is shown as a function of time and aligned to the offset of the fixation point that served as a command to the NHP to respond (thin vertical line at time 0 ms).

**Figure 2:**
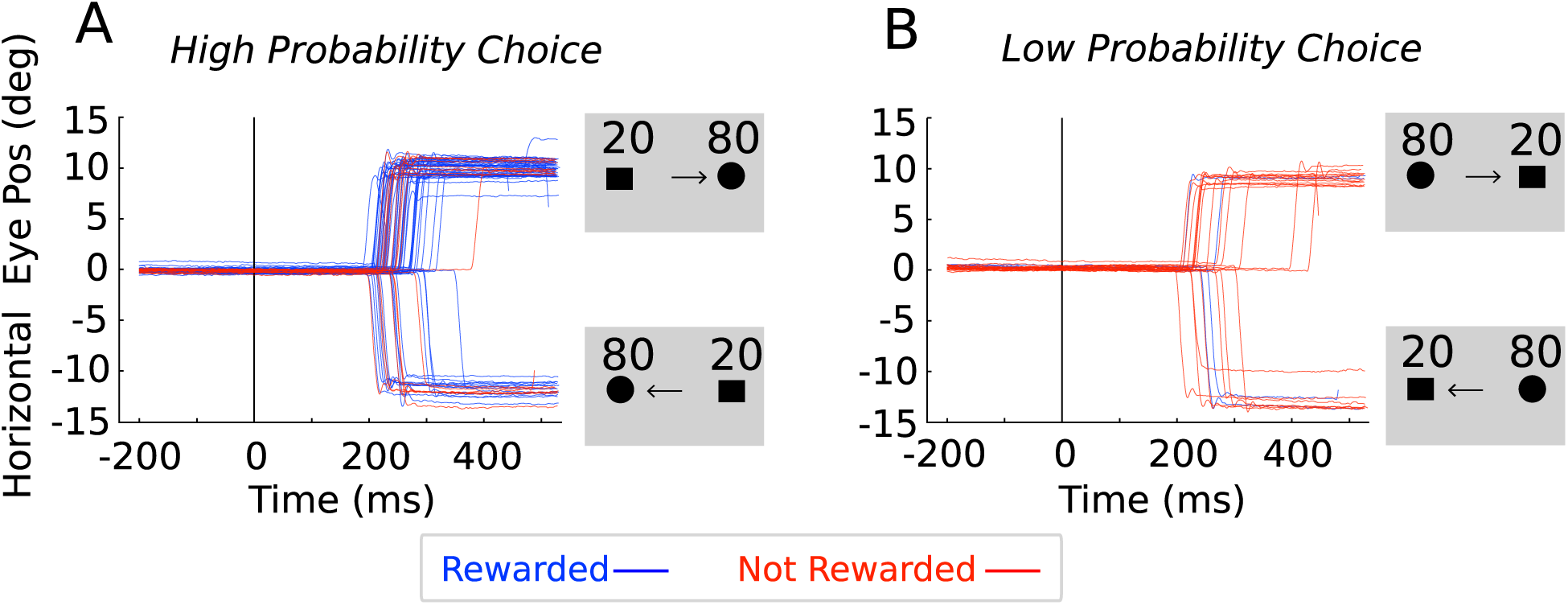
Representative horizontal component of eye movements from an nonhuman primate (NHP) performing the probability switching task (PST) during a task-based functional magnetic resonance (fMRI) session. In the block of trials illustrated in this example, the circle target is assigned an 80% chance of reward, with the square receiving a reward 20% of the time. The end points of the saccadic eye movements are used to assess the NHP’s choices. (A) High probability target selections, and (B) low probability target selections. Rewarded trials are shown in blue, and non-rewarded trials are shown in red. A representation of the targets, their corresponding probability assignments, and arrows indicating the NHP’s choices, are illustrated in the four gray insets.

Trials corresponding to the high reward probability (*correct*) choice are shown in Figure 2A, and those corresponding to the low probability choice are shown in Figure 2B, with rewarded trials drawn in blue and non-rewarded trials drawn in red. For the blocks of trials shown, the circle target is the correct choice. However, this correct choice is rewarded on only 80% of the trials. In contrast, the incorrect choice is rewarded on 20% of the trials.

The data were processed with DataViewer (SR Research, Ottawa, Canada). For each trial recorded in each experimental session, the target locations and probabilities, subject choice, and reward were exported for analysis. Over multiple blocks, we computed the probability of correct response for each of the 20 trials before and after the switch of probabilities.

NHP GD performed in three different probability schemes: 100*/*0, 90*/*10, and 80*/*20; the total numbers of switches for these schemes were 44, 31, and 49, respectively. NHPs M and G only performed in the 80*/*20 probability scheme; the total number of switches was 5 and 25, respectively.

### Deep Recurrent Neural Network Model

We present an abstract model of sequential decision making that uses only action choice for a given trial and the resulting, potentially stochastic, outcome signal to prepare to act in the subsequent trial. Our goals are to provide computational accounts for 1) what information must be represented from one trial to the next (i.e., the belief state), 2) how action choices are made given this information, and 3) the behavior that results under different reward probability schemes. Furthermore, we do not impose a specific rule for updating the model’s belief state, but instead allow the model to choose the details of this update.

We present the model with an abstract version of the PST task. This task is naturally expressed as a Partially Observable Markov Decision Process (POMDP) because it requires the agent to perform a sequence of actions for which the outcome of individual trials is ambiguous (Boutilier and Poole, 1996; Kurniawati, 2022). Specifically, there exists a true trial state that encodes the preferred action and the probability of reward. However, the NHP and model agents only have access to trial outcome in the form of a reward. When this reward is stochastic, the outcome for a single trial is ambiguous as to the true trial state. This requires the agent to integrate action and outcome information across multiple trials to form a richer belief state upon which “ideal” action decisions can be made. Complete solutions to POMDPs are computationally intractable, but efficient solutions can be approximated using recurrent neural networks (RNNs) to implement belief state estimation, and a Q-function to estimate the value of taking an action given the estimated state (Boutilier and Poole, 1996; Rao, 2010; Huang and Rao, 2013; Lambrechts et al., 2022a; Rao, 2024).

Each task session is composed of two blocks of trials. For each trial, the model agent must use its estimate of belief state to select one of two actions (*A*_0_ and *A*_1_), which are analogous to the saccade to square/circle choices made by the NHPs. Within the first block of trials for session *s*, *A*_0_ is rewarded with probability of *p_s_* and *A*_1_ is rewarded with probability 1 − *p_s_*. After a randomly chosen number of trials, the task switches the reward relationship: 1 − *p_s_* and *p_s_* for actions *A*_0_ and *A*_1_, respectively. As with the NHP task, the model agent is not cued as to the base session probability nor the time of switch.

The objective of the model training process is for the agent to identify a rule for selecting actions that maximizes the expected accumulated reward into the future. More specifically, for a given trial *t*, the goal of the model is to maximize:

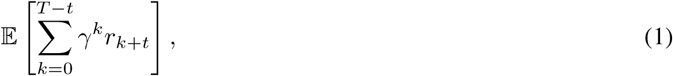

where 0 *< γ <* 1 is a discount factor for the value of future rewards, and T is the total number of trials in the session. Following a DRQL modeling approach, the belief state estimation and subsequent action choice for a single trial are captured as one model step. The components that implement this model step for trial *t* are shown in Figure 3. The belief state at trial *t*, *X_t_* ∈ R*^N^*, can be approximated as a function of the information available from the previous trial:

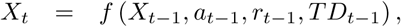

**Figure 3:**
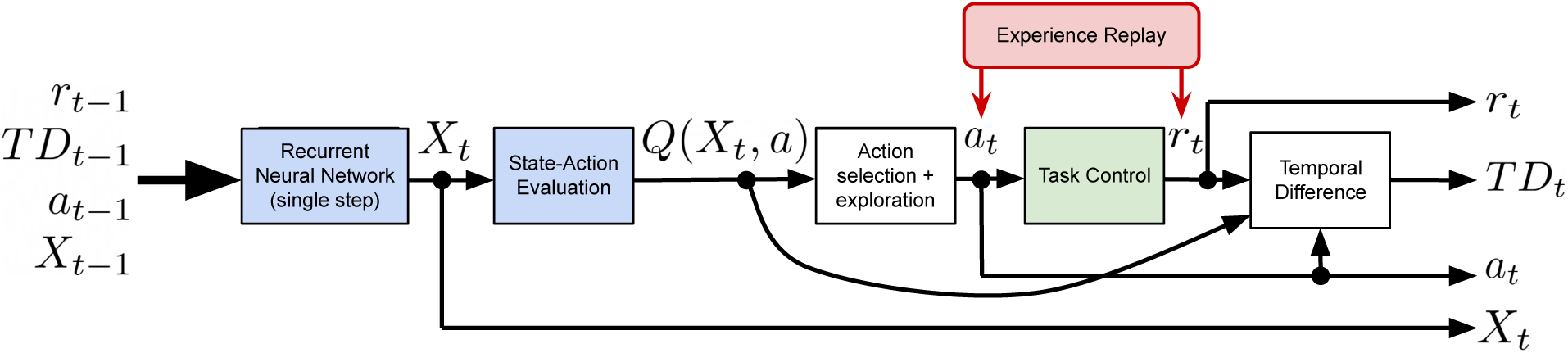
Single step of the model architecture. Agent components are shown in white (fixed logic) and blue (learned logic); the task component is shown in green. *X_t_* is a vector of N modeled neurons at time *t*; *Q*(*X_t_, a*) represents the evaluation of the combination of the current state and each possible action; *a_t_* is the selected action to be executed; *r_t_* is the reward that is given in response to the selected action; and *TD_t_* represents the temporal difference error between the expected outcome of an action and the actual outcome. Under evaluation conditions, the actions and resulting rewards from a trained model may be substituted for the behavior and consequences of another agent (NHP or another model; red box).

where *X_t_*_−1_ ∈ R*^N^* is the previous belief state, *a_t_*_−1_ ∈ {*A*_0_*, A*_1_} is the last executed action, *r_t_*_−1_ ∈ {0, 1} is the last received reward, and *TD_t_*_−1_ ∈ R is the model’s estimated temporal difference error (defined below). *f* (.) is a non-linear function implemented as a feed-forward neural network with trainable parameters.

The state-action value, *Q*(*X_t_, a*), represents the expectation of the future rewards, as defined in Equation 1, assuming that action *a* is taken from belief state *X_t_*, followed by a sequence of actions selected by a fixed behavioral rule (Watkins and Dayan, 1992; Hausknecht and Stone, 2015). Because the true Q-function is unknown, we estimate it using a non-linear function for each action:

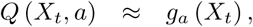

where *g_a_*(.) is a trainable function for action *a* that is also implemented as a fully-connected neural network with one output unit for each action.

Under ideal conditions, the best performing behavioral rule is one that *greedily* chooses the action at trial *t*:

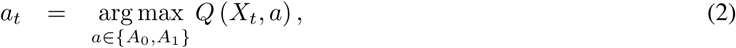

so long that *Q*(*X_t_, a*) also reflects this same behavioral rule. During the training process, a certain degree of action exploration is necessary to ensure that both action choices are considered from a sufficient distribution of belief states. Here, we take a simple (and standard) *ɛ*-greedy approach to action choice, in which the agent chooses a random action with probability *ɛ*, and otherwise chooses the greedy action, as defined in Equation 2.

Based on this chosen action and the true (but unknown to the agent) task state, the model agent is assigned a reward, *r_t_*, as described above. Finally, the agent computes a *temporal difference (TD) error*, which is a measure of the difference between the agent’s expectation of value of executing action *a_t_*_−1_ on the previous trial, *Q* (*X_t_*_−1_*, a_t_*_−1_), and the actual outcome of executing that action. The latter term is a sum of the received reward, *r_t_*, and the agent’s estimate of the best it can do from the subsequent state. Specifically:

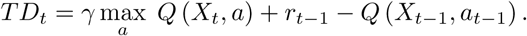

The best performing behavioral rule (again, under ideal conditions) will minimize the expected value of the TD error over all possible sessions. With DRQL we iteratively measure TD error over a sample of these sessions (S):

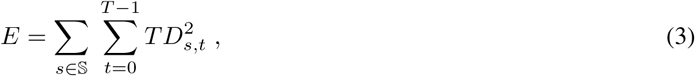

and adjust the parameters of *f* (.) and *g_a_*(.) to reduce *E*.

### Model Training Protocol

A model session is composed of two blocks of trials, consisting of a total of *T* = 200 trials. The initial state of the model is initialized to be neutral (*r* = 0, *TD* = 0, *a* = [0, 0], and *X* = [0, 0*, …*0]). The switch between the two blocks is randomly selected on trial 20 ≤ *t < T* − 20. In addition, the baseline reward probability is selected randomly to be one of 100%, 90%, 80%, 70%, or 60%, and the action with the highest reward probability is selected randomly to be either *A*_0_ or *A*_1_. These choices yield 1600 different session configurations, from which 1024 are selected with replacement for each of a training data set and a separate validation data set. The expected number of session configurations in common between the training and validations data sets is ≈ 358*/*1024 sessions.

We perform model training in parallel: the current model is presented with each session in the training data set, from which *E* is measured with respect to the training sessions. We then perform a gradient descent step. This process is repeated until *E* (measured with respect to the validation sessions) does not improve substantially over a sequence of training epochs. Following this training, the model parameters are frozen for any subsequent analysis. This trained model can then be presented with any session configuration. For each trial within the session, the model updates its recurrent state and Q-value estimates, from which the Q-optimal action is identified and an *ɛ*-greedy action is chosen for execution (Figure 3). The Task Control module then assigns a reward and the model computes a TD error.

### Model Implementation Details

#### Single Trial Module

The model is implemented using the Keras 3 (Chollet and the Keras 3 Development Team, 2015) and Tensorflow 2.18 (Abadi et al., 2015) packages for Python 3.12 (van Rossum and de Boer, 1991). Figure 3 summarizes the implementation of the computational processes for a single trial; the state variables are listed in Table 1. The two plastic components are the recurrent neural network that computes the update to the belief state (*f* (.)) and the feed-forward neural networks that estimate the Q-values for each action (*g_a_*(.)). The structural details of these neural networks are shown in Table 2.

**Table 1:** Key model variables.

|  | Variable | Dimensions | Notes |
| --- | --- | --- | --- |
| Reward | $r$ | 1 | |
| Temporal Difference Error | $TD$ | 1 | |
| Executed action (RNN input) | $a$ | Number of actions (2) | one-hot encoding |
| Executed action (selected action) | $a$ | 1 | integer |
| Belief state | $X$ | 10 | |
| Q-values | Q | Number of actions (2) |  |

**Table 2:** Neural network configuration.

|  | Recurrent Neural Network | Q-Value |
| --- | --- | --- |
| | $f(\cdot)$ | $g_a(\cdot)$ |
| # Hidden neurons | n/a | 10 |
| Hidden non-linearity | n/a | exponential linear unit (elu) |
| # Output neurons | 10 | 1 per action (2) |
| Output non-linearity | hyperbolic tangent (tanh) | linear |
| Total # of parameters | 150 | 132 |

The remaining components of the single-trial module (Action selection + exploration, Task Control, and Temporal Difference) are implemented as fixed computational components within Keras 3.

#### Full Model

The full *N* = 200 trial model is implemented as an *unrolled* recurrent neural network model. Multiple virtual copies of the single trial module are wired back-to-back, with the state variables that serve as outputs of trial *t* providing the inputs to trial *t* + 1. However, the adjustable parameters within *f* (.) and *g_a_*(.) are shared across the different copies.

Keras 3 and Tensorflow automatically provide the computational machinery for backpropagating errors through the full model. Specifically, squared TD error across each step of the model can influence the error computation in prior trials’ Q-values, belief state, and associated parameters (see Equation 3). Although the *Task Control* module is implemented as part of the Keras 3 model, error gradients are disallowed from flowing through this computation.

#### Model Training

The model training choices, including hyper-parameters are shown in Table 3.

**Table 3:** High-level model choices.

| Hyper-Parameter / Model Choice | Value |
| --- | --- |
| Loss function | $\sum TD^2$ |
| Exploration fraction ($\epsilon$) | 0.1 |
| Loss function | $\sum TD^2$ |
| Regularization | n/a |
| Optimizer | Adam |
| Training epochs | 10,000 |
| Early stopping patience | 2,000 epochs |
| Early stopping monitor variable | training loss |
| Early stopping min delta | $10^{-4}$ |
| Learning rate | $10^{-3}$ |
| Training set size | 1,024 sessions |

### Experience Replay

*Experience replay* (ER) is the process of presenting a RL model with a prerecorded sequence of actions and outcomes, overriding the model’s own action choices (and subsequent outcomes), but still computing belief state and the associated Q-values (Lin, 1991; Roscow et al., 2021). Typically, ER is used to allow training of the model based on experiences produced by another agent or by the same learning agent, but collected at some other time. Here, we use ER not as an approach for training, but as one to examine the sequence of belief states and Q-values as if the model were following the same action/outcome trajectory of some other agent. When the other agent uses a different trained model, this gives us a basis for comparing the consistency of the models’ task representations. However, when a NHP is the other agent, the ER approach provides a hypothesis as to the types of information that must be represented by the central nervous system as the NHP is performing the switching task. When the NHPs are performing blocks of trials, it is not uncommon for the NHP to fail to perform a left or right saccade action. Because the current model does not represent *non-performance* responses, we remove these trials for the purposes of the ER analysis.

### Statistical Analysis

We measure model and NHP performance in a task with respect to the frequency of their selection of the action with the highest probability of reward. For both the deterministic and the probabilistic tasks, we refer to this measure as *percent correct*.

When comparing the Q-values produced by two models in the same task and under the same behavioral / outcome sequence, we report the *Fraction of Variance Accounted For* (FVAF) by one model with respect to the other. For a corresponding set of scalar samples (*y_i_* and *x_i_*, *i* ∈ {0*…M* − 1}), FVAF is defined as:

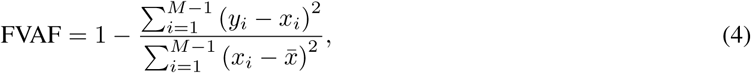

where *M* is the number of samples, the numerator is the sum squared error between *y* and *x*, and the denominator is a factor of the variance of *x*. FVAF is unit-less and has a similar interpretation as *R*^2^. FVAF = 1 represents a perfect equivalence of *y* with *x*; smaller positive values represent a reduction of this alignment; FVAF = 0 means that there is no alignment of the two; and FVAF *<* 0 implies no alignment with the squared error having a higher variance than *x*.

We perform hypothesis testing using a two-tailed student t-test.

### Software

Data handling and figure generation are performed using Pandas 2.3 (Pandas Development Team, 2025) and Matplotlib 3.10 (Hunter, 2007), respectively.

### Code Accessibility

All code is available at Zenodo: https://doi.org/10.5281/zenodo.18926284

### Data Accessbility

All trained models and resulting data are available at Zenodo: https://doi.org/10.5281/ `zenodo.18932110`

## Results

The probability of receiving a reward alters the behavior of both the NHPs and the model agent. Figure 4A shows the behavior of one NHP (GD) for 200 trials under the 100*/*0 probability scheme (i.e., a deterministic task). The task (dark blue curve) is to perform action *A*_1_ for the first 88 trials, switching to *A*_0_ at trial 89, and then switching back to *A*_1_ at trial 174. The NHP performs *A*_1_ until trial 91 (gray). Because the task switched at 89, the response is considered *not correct* for trials 89 − 91 (light blue); furthermore, because this is a deterministic task, the reward delivered to the NHP (olive) exactly tracks the correctness curve. While the task is to produce action *A*_0_, the NHP largely does so, but occasionally explores with *A*_1_. This exploration is met with no reward. After the task switches back to *A*_1_ at trial 174, the NHP continues to produce *A*_0_ until trial 182. The incorrect response during this time is met with no reward.

**Figure 4:**
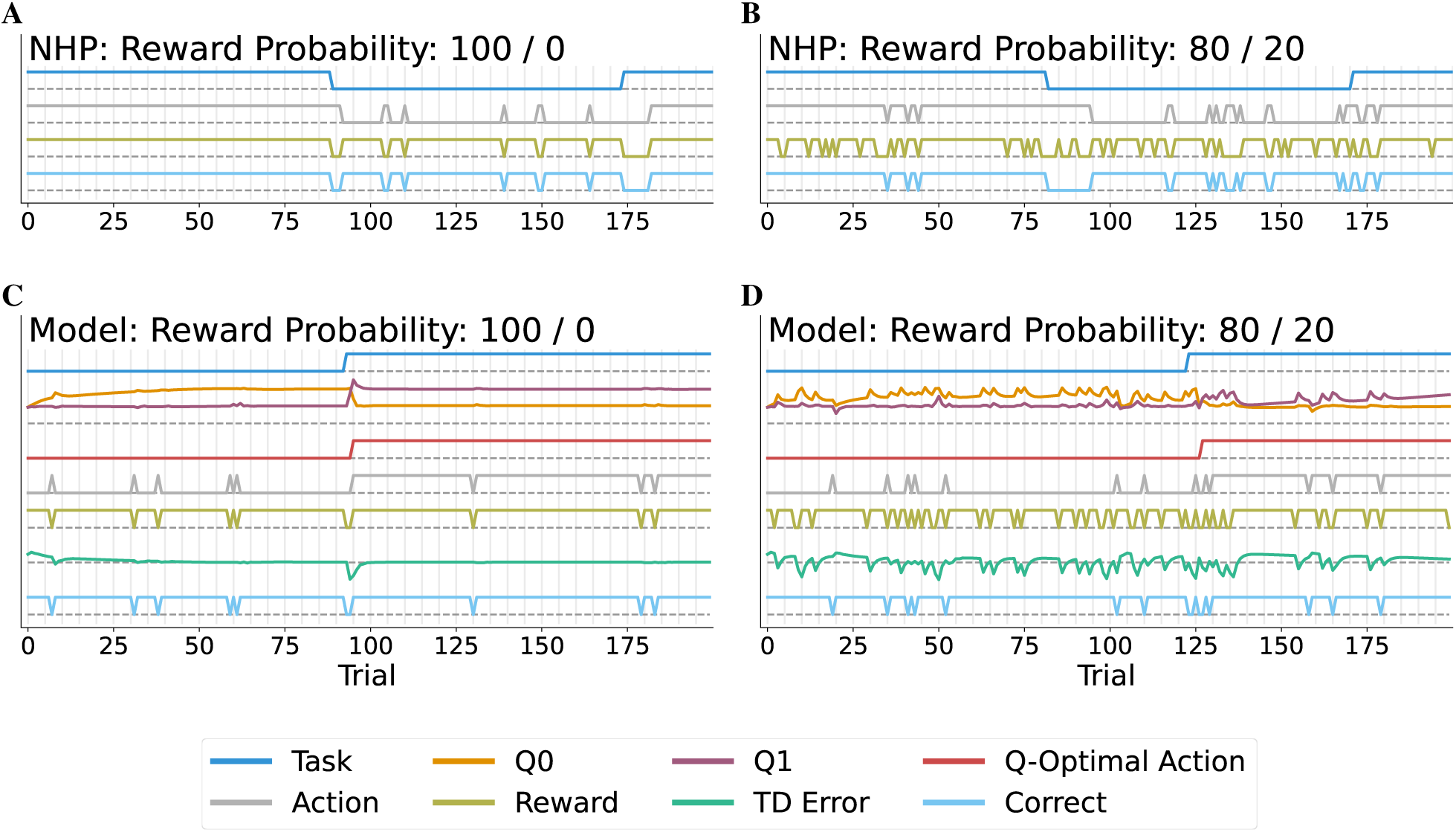
Behavior during 200-trial sessions involving at least one task switch for a NHP (A and B) and for the trained model (C and D). In panels A and C, actions are rewarded with probability 100% when they match the current task, and 0% when they do not; in panels B and D, the actions are rewarded with probability 80% and 20%, respectively. The current task (dark blue) is not explicitly cued to the NHP/model agent. The action chosen by the agent is shown in gray, the resulting reward is shown in olive, and whether the response correctly matched the task is shown in light blue. The latent variables within the model (C and D) are the Q-values for actions *A*_0_ (orange) and *A*_1_ (purple), the Q-optimal action (red), and the TD error (blue-green).

Under the 80*/*20 probability scheme, the high probability action is only rewarded 80% of the time, and the low probability action is rewarded 20% of the time. Figure 4B shows the NHP response during such a block of 200 trials. The task is to produce action *A*_1_ for the first 81 trials. During this time, the NHP explores with action *A*_0_ on three trials. The first and last of these exploratory actions went unrewarded, but the middle action at trial 41 was rewarded. After the task is switched to *A*_0_ at trial 82, the NHP continues to produce *A*_1_ until trial 95, receiving a reward for three of the incorrect responses. During this middle period between switches, the NHP continues to explore by producing *A*_1_. In fact, the NHP produces *A*_1_ on the very first trial that the task is switched to requiring action *A*_1_ at trial 171, and is subsequently rewarded. However, when the NHP repeats *A*_1_ on trial 172, it is unrewarded. The NHP responds by exploring both actions before settling on *A*_1_ at trial 179.

With the model, in addition to the behavioral response and subsequent reward, we are able to inspect variables that are internal to the model (Figure 4C,D), including the Q-values (*Q*_0_ = orange; *Q*_1_ = purple), the action with the highest Q-value (the *Q-optimal action*; red), and the temporal difference error (blue-green). Under the deterministic task (panel 4C), the Q-values are initially similar values, but with a few repeated correct *A*_0_ actions, *Q*_0_ gradually increases to a higher value. During the first 92 trials, the model does explore occasionally; this is due to the fixed exploration probability of 10%. These exploratory actions go unrewarded; however, the model anticipates these non-rewards, as seen in the small variations in the TD Error. When the task switches at trial 93, the model responds with *A*_0_ since it has no cues as to the switch. On trial 94, it begins to integrate the prior non-reward into its belief state, as shown by the changes in the Q-values. By the second incorrect *A*_0_, which is also not rewarded, the Q-values cross each other, switching the Q-optimal (and selected) action to *A*_1_ on trial 95. During this transition, the TD error also drops, indicating a transient unanticipated (and negative) expectation of future reward. Note that this change is delayed by one trial because the TD error computation relies on the Q-values associated with both the current *and next* belief states.

When faced with the 80*/*20 task, there are several key changes to the model’s latent representation and its behavior when compared to the deterministic case (Figure 4D). First, the average difference between the two Q-values is smaller. This captures the greater uncertainty in interpreting an *apparently correct* action that is unrewarded – it could be due to a switch in task (so, the action was incorrect) or it could be that the action was correct. In addition, the TD error often deviates from zero during the block of trials; this is due to the unexpected non-rewards for correct actions (or reward for incorrect actions). The task actually switches to *A*_1_ at trial 123, but the Q-values do not cross until trial 127. Although *A*_1_ at this trial is the Q-optimal action, the model chooses to perform an exploratory *A*_0_, which is unrewarded. It is not until trial 130 that the model begins to consistently perform *A*_1_. However, due to several immediate non-rewards, *Q*_1_ drops and requires the remaining trials to fully recover. This may be due to the model considering the hypothesis that it is actually performing a task with lower probability of reward for correct actions.

### Average Behavior Around Task Switch

The response of the model and the NHP to a task switch varies depending on the probability of reward, as shown in Figure 5. Across all blocks, we aligned the trials that the switch occurred and computed the average correctness relative to this time of switch (relative trial 0 is the first trial of the new action). Figure 5A shows the average correctness for the model across 1024 task switches, split across the five different probability schemes. The baseline correctness prior to the switch for tasks 100*/*0, 90*/*10, 80*/*20, and 70*/*30 hovers around 90%. This corresponds to the fixed 10% probability of exploration on any given trial. Task 60*/*40 is degraded from this baseline, likely due to higher uncertainty as to when the switch has occurred. Following the switch, the model performs very poorly on the next two trials (0 and 1). For the 100*/*0 task, the model crosses 50% correctness and nearly recovers performance by trial 2; and it achieves baseline performance by trial 3. However, the non-deterministic tasks require more time to recover, with 90*/*10, 80*/*20, 70*/*30, and 60*/*40 crossing 50% around trials 3, 4, 8, and 10, respectively. While 90*/*10 and 80*/*20 return to baseline by trial 9, 70*/*30 and 60*/*40 are still recovering by trial 19 after the switch.

**Figure 5:**
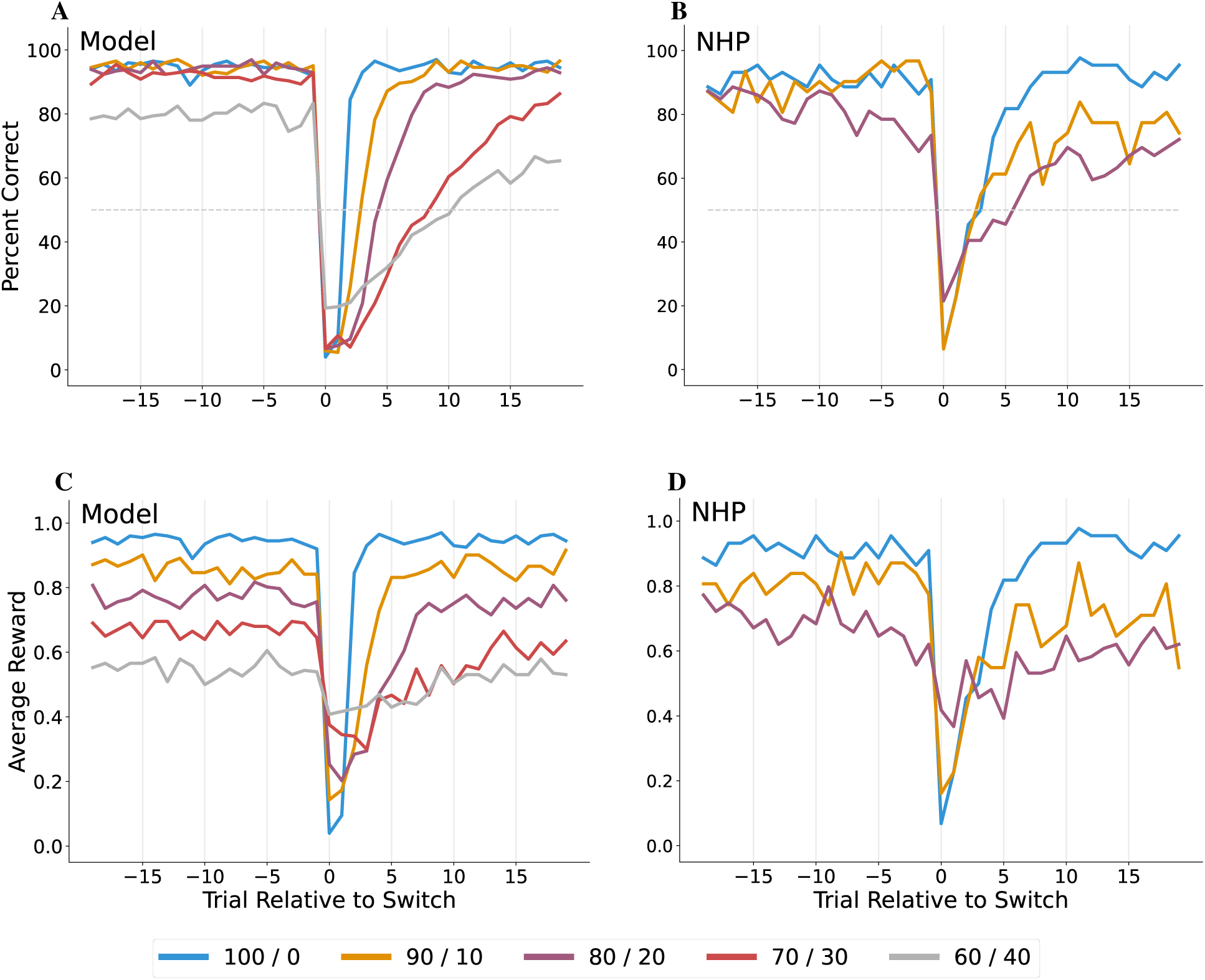
Percent of correct responses relative to the time of switch from one task to the other for the model (A) and the NHP (B); average reward relative to the time of switch for the model (C) and the NHPs (D). Trial 0 is the first trial after the switch. Each curve represents the different reward probability conditions: 100*/*0 (blue), 90*/*10 (orange), 80/20 (purple), 70*/*30 (red), and 60*/*40 (gray). The NHPs are only presented with the first three conditions.

The NHPs exhibit a similar recovery behavior. Figure 5B shows the average correctness over a total of 154 switches across three NHPs. On average, the deterministic case requires approximately 8 trials after the switch to return to baseline. In contrast, 90*/*10 and 80*/*20 are still returning to baseline by trial 19, though the latter returns at a slower rate.

Figure 5C,D shows the average reward around the time of switch, separated by probability scheme for the model and the NHP, respectively. For the model, the average reward between neighboring probability schemes is separated by approximately 0.1. In contrast, for the NHPs, the average reward for each scheme is lower than the model, and the difference between neighboring schemes is larger than 0.1. These differences likely reflect the NHP’s tendency to explore more (even in the deterministic case) and to increase the probability of exploration as the reward becomes more uncertain.

### Experience Replay

While the trained model is capable of forming an internal representation of the task from prior trial feedback and then making appropriate action choices, it is possible to replace the model’s action (and resulting reward) with the action choice made by the NHP (and the reward given to the NHP). This allows the model to form a latent representation of the task as performed by the NHP, which may be suggestive of the types of information that must be encoded in the NHP’s central nervous system. Figure 6 shows the result of replaying the NHP behavior from a 100*/*0 trial block (A) and a 80*/*20 trial block (B) into the model (the same trial blocks from Figure 4A,B). For the deterministic block, the task switches at trial 89. The Q-values cross immediately at trial 90. However, the NHP still chooses action *A*_1_ through trial 91. After the switch, the NHP explores with several *A*_1_ actions, receiving no reward. Nonetheless, the model TD error does not change substantially from zero, implying that the model expected the non-reward. The task switches back to *A*_1_ at trial 173, and the Q-values cross at trial 176. However, the NHP does not switch to *A*_1_ until trial 182.

**Figure 6:**
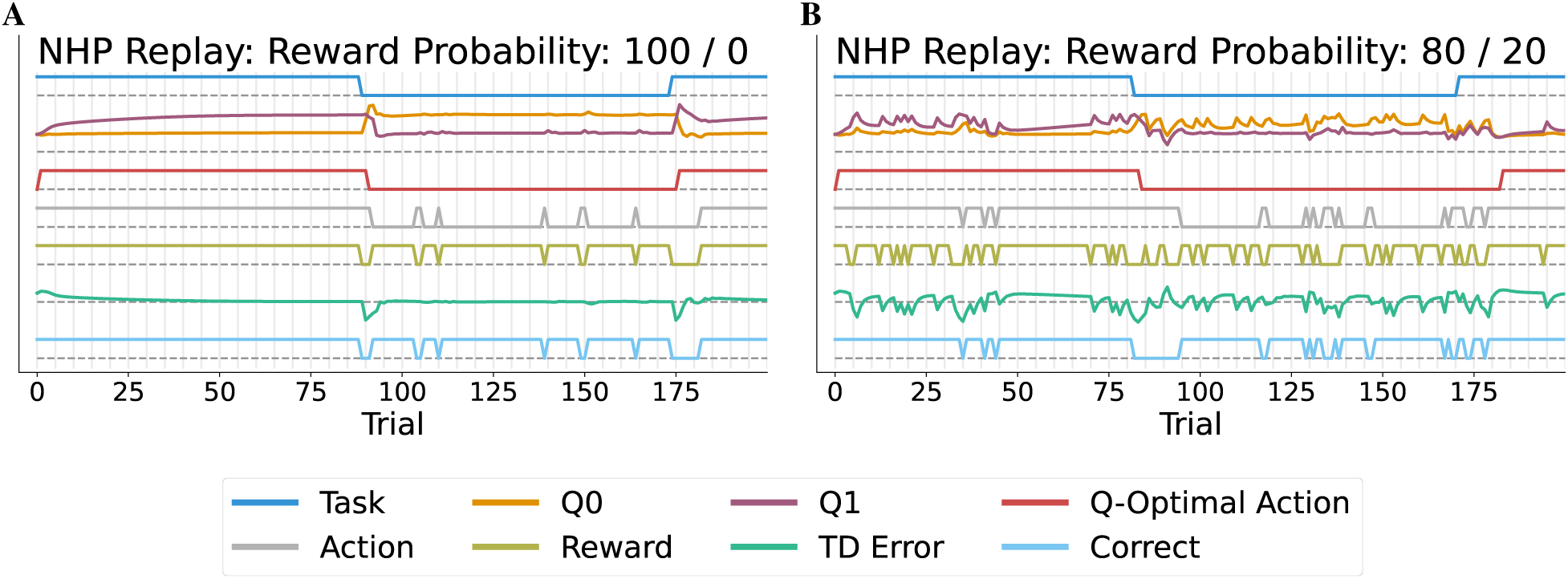
Model latent variables when the chosen action is determined by the NHP behavior and the reward is chosen in response to the NHP for the 100/0 (A) and 80/20 (B) reward conditions. The notation is the same as in Figure 4.

For the 80*/*20 probability scheme (Figure 6B), much like when the model makes its own action choices, the Q-values on average differ less than in the deterministic case. Yet, after the task switch at trial 81, the model does not recognize the switch until trial 84, and the NHP does not switch until trial 95. In contrast, for the task switch at trial 171, the model does not recognize the switch until trial 183. Although the NHP does switch earlier, it does not produce a consistent action until trial 179. In part, this delay appears to be due to rewards that are inconsistent with the task (a non-reward for a correct action at trial 172 and a reward for an incorrect action at trial 177).

### Belief States and Q-Values

The trained model allows for the examination of its internal latent variables around the time of a switch in task. Figure 7A,C shows the average belief state for two selected recurrent neurons (the remaining recurrent neurons are shown in Figures S2A,C,E, S3A,C,E, and S4A,C). For these figures, the 1024 trial blocks for the model have been grouped by probability scheme (color) *and* initial expected action (*A*_0_: solid, and *A*_1_: dashed), aligned at the time of switch, and then averaged by group. The dominant variation of recurrent neuron 0 appears to be with respect to the probability scheme, with the deterministic task being encoded with higher activation levels, and the probabilistic tasks being encoded with progressively lower activation levels. This recurrent neuron responds to the time of switch; the degree of change at the switch may encode the amount of “surprise” that the model has for the unexpected non-reward. Note that change in modulation does not occur until one trial after the switch; this is due to the fact that the model is not “aware” of the switch until it observes the outcome of relative trial zero. In addition, by trial 19 after the switch, this neuron has nearly returned to its baseline prior to the switch. However, for the non-deterministic cases, the relative degree of modulation between *A*_0_ first (solid) and *A*_1_ first (dashed) have flipped. In contrast, prior to the switch, recurrent neuron 9 does not make a substantial distinction between probability schemes. Instead, this neuron distinguishes the appropriate action. Following the switch, this neuron flips polarity, with the rate of flip being dependent on the probability of reward.

**Figure 7:**
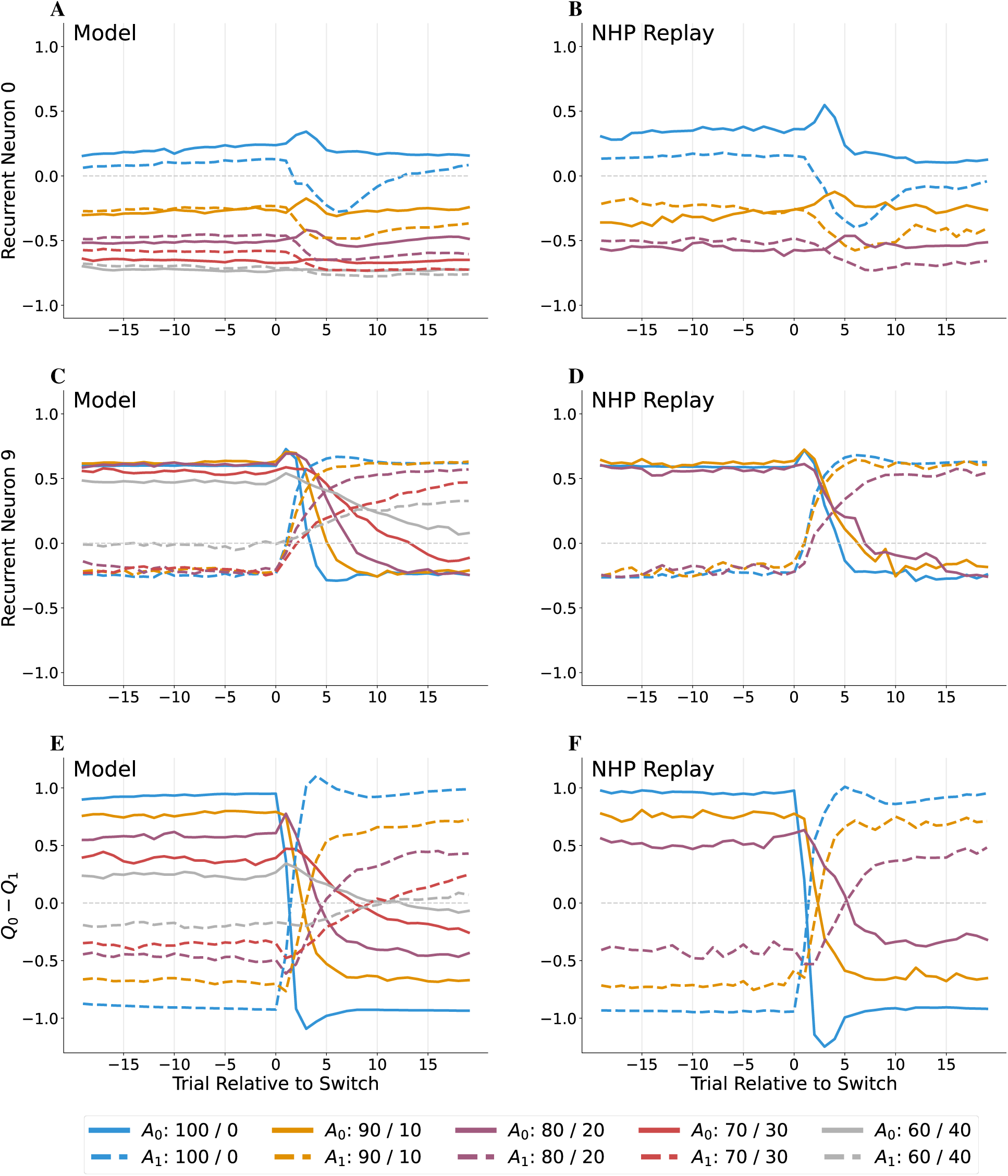
Average model belief state variable response relative to the time of switch for different reward conditions (color) and starting action (*A*_0_=solid; *A*_1_=dashed). Model neuron 0 and 9 when the model determines the actions (A and C) and when the NHP GD behavior is replayed into the model (B and D). Difference between the Q-values for actions *A*_0_ and *A*_1_ when the model determines the actions (E) and when the NHP behavior is replayed (F).

Figure 7B,D show the average activity of the same neurons when the NHP GD behavior is replayed into the model.

These neurons show very similar behavior to when the model is making action decisions. However, the switch of polarity of neuron 9 (from positive to negative, and vice-versa) is delayed by several trials for the NHP replay case, which is consistent with the NHP requiring more time to switch than the model.

In addition to belief state, it is possible to examine the average difference in Q-values near the task switch (Figure 7E,F for the model and NHP replay cases, respectively). As in the other panels, each curve represents the average difference, separated by probability scheme and initial action. A positive difference corresponds to action *A*_0_ being more favored than action *A*_1_. Prior to the switch and when the task is to produce *A*_0_, this difference is positive (solid), indicating that the model has a high confidence of receiving, on average, reward in future actions. When the task is deterministic (blue), the magnitude of the Q-value difference is highest because the expectation is that subsequent trials will be rewarded consistently. As the probability of reward is reduced (other colors), the expectation of future rewards becomes lower, as encoded by the smaller difference between the Q-values. On the contrary, when the initial task is to generate *A*_1_, the Q-value difference is negative (dashed), indicating that action *A*_1_ is preferred leading up to the switch.

Following the task switch, the curves move toward the opposite sign. When a given curve crosses zero, this corresponds to the trial at which the Q-optimal action switches from one action to the other. For the model (panel E), there are two features of note. First, each *A*_0_-to-*A*_1_ zero crossing occurs at the same point as the *A*_1_-to-*A*_0_ crossing. This is indicative of a symmetry in the over-trained model. Second, the change in Q-optimal action on average takes place faster for the deterministic task; this switch takes longer for progressively lower probability tasks.

When the NHP behavior is replayed into the model (Figure 7F), the Q-value difference curves follow a similar time-course as when the model agent is making action decisions. In particular, the zero-crossing of the differences occur at approximately the same point in time. This contrasts with the NHP’s delayed recovery after the switch in producing the correct action (compare Figure 5B vs. A).

### Temporal Difference Error

Temporal difference error captures the difference between the expected value of taking a specific action and the actual outcome of taking that action. Recall that values encode not only the immediate reward, but also the expectation of future rewards if the Q-optimal action is consistently selected thereafter. Figure 8 shows the TD error for each probability scheme relative to the time of switch for the model (A) and for the NHP behavior replayed into the model (B). TD errors prior to the switch are approximately zero, implying that the model is anticipating the rewards on average for any of the schemes. The exception in panel A is the 60*/*40 scheme, in which the TD error is around −0.1, indicating that the model is overly optimistic in its expectation of future rewards.

**Figure 8:**
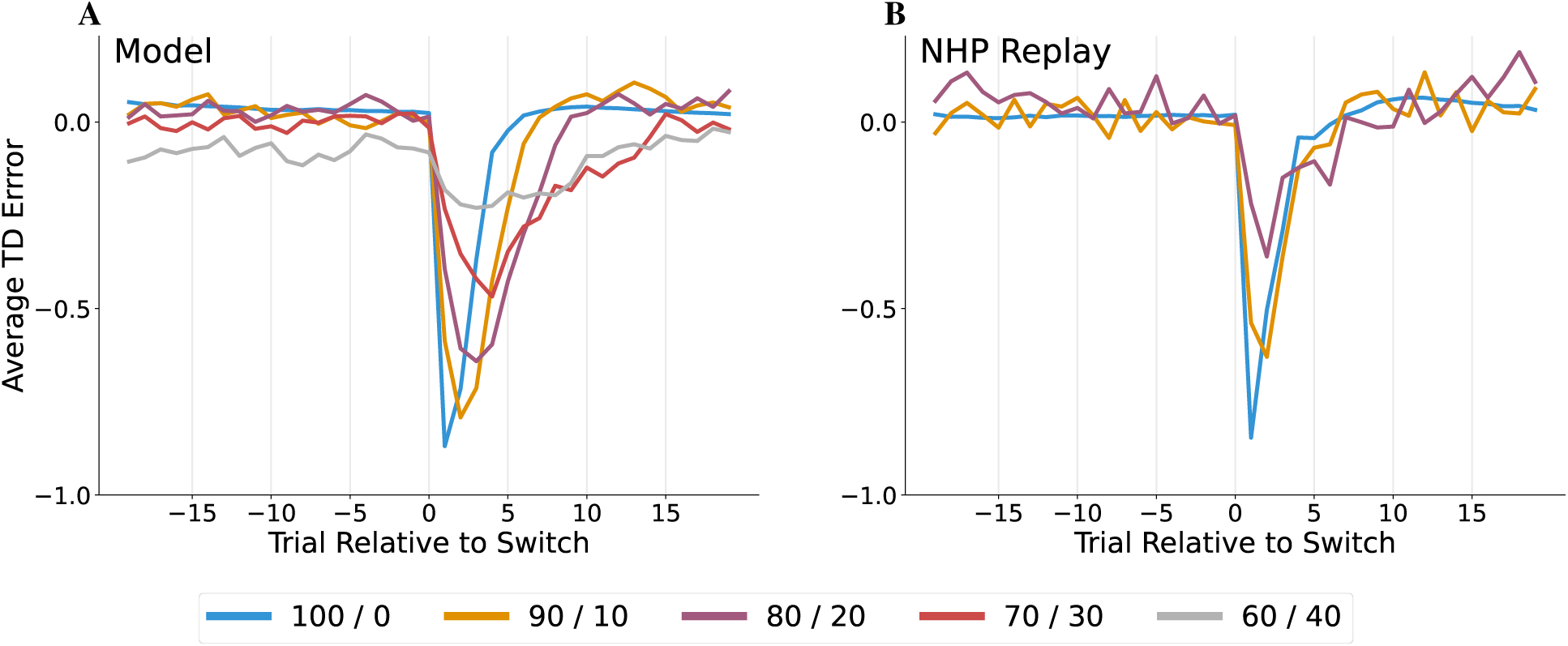
Average TD Error relative to the time of task switch when the model selects the actions (A), and when the NHP GD behavior is replayed into the model (B).

After the switch, for the deterministic case, the magnitude of the TD error is largest, with the magnitude becoming smaller as the probability of reward drops. This effect is due to two factors. First, the Q-value for the executed action is larger for the higher probability cases (capturing the expectation of future rewards if the task had not switched). After the switch, continuing to select this action means that a corresponding reward is unlikely, which violates this expectation. Second, following the switch, continuing to select the original Q-optimal action will still garner some reward; in fact, the probability of reward in this case is highest for the low probability schemes (e.g., 40% for the 60*/*40 scheme). So, on average, violation of the expectation is smaller in magnitude. Also note that the time to recovery for the TD error increases with less certainty in the reward. This is due to the need to accumulate more trial outcome observations before the model can commit to the switch.

When the NHP behavior is replayed into the model (Figure 8B), average TD error for the first three probability schemes follow a similar time-course, both in magnitude and timing, as when the model is selecting actions. However, it is unclear whether there are differences in the time to recovery across the three probability schemes. This may be due to differences in exploration strategy, as well as the smaller number of switches that are included in the TD error averages.

### Belief State Information Content

Figure 7A–D shows that the behavior of recurrent neurons can capture different aspects of the overall task (namely, the probability of reward, and which action should be generated). However, what key information is captured in the belief state, as encoded by the full set of recurrent neurons? One approach is to project the 10-dimensional recurrent state into the first two principal components of variation of this recurrent neuron set. While the PCA model is computed using all time steps and all trial blocks in the training data set, the first two components of model capture 81% of the variance observed in the full validation data set. Figure 9 shows the average projection into the space defined by the first two principal components around the time of switch for the model (A,C) and for when the NHP GD behavior is replayed into the model (B,D). Panels A,B represent the transition from *A*_0_ to *A*_1_, where panels C,D show the opposite transition. Each point is the average projected coordinate for a specific trial relative to the switch, and each color represents a different probability scheme. The point centered at the diamond is the average projection at the trial of switch (but before the model has access to the outcome of the action taken at the switch). For the trials leading up to and including the switch, the model has a consistent average belief state representation, as indicated by the cluster of points in the neighborhood around the diamond. However, after the model begins to incorporate the post-switch outcome information, the average projection begins to shift to the left for the *A*_0_ → *A*_1_ transition (panels A,B), and right for the *A*_1_ → *A*_0_ transition (C,D). Trial 19 after the switch is marked with a circle.

**Figure 9:**
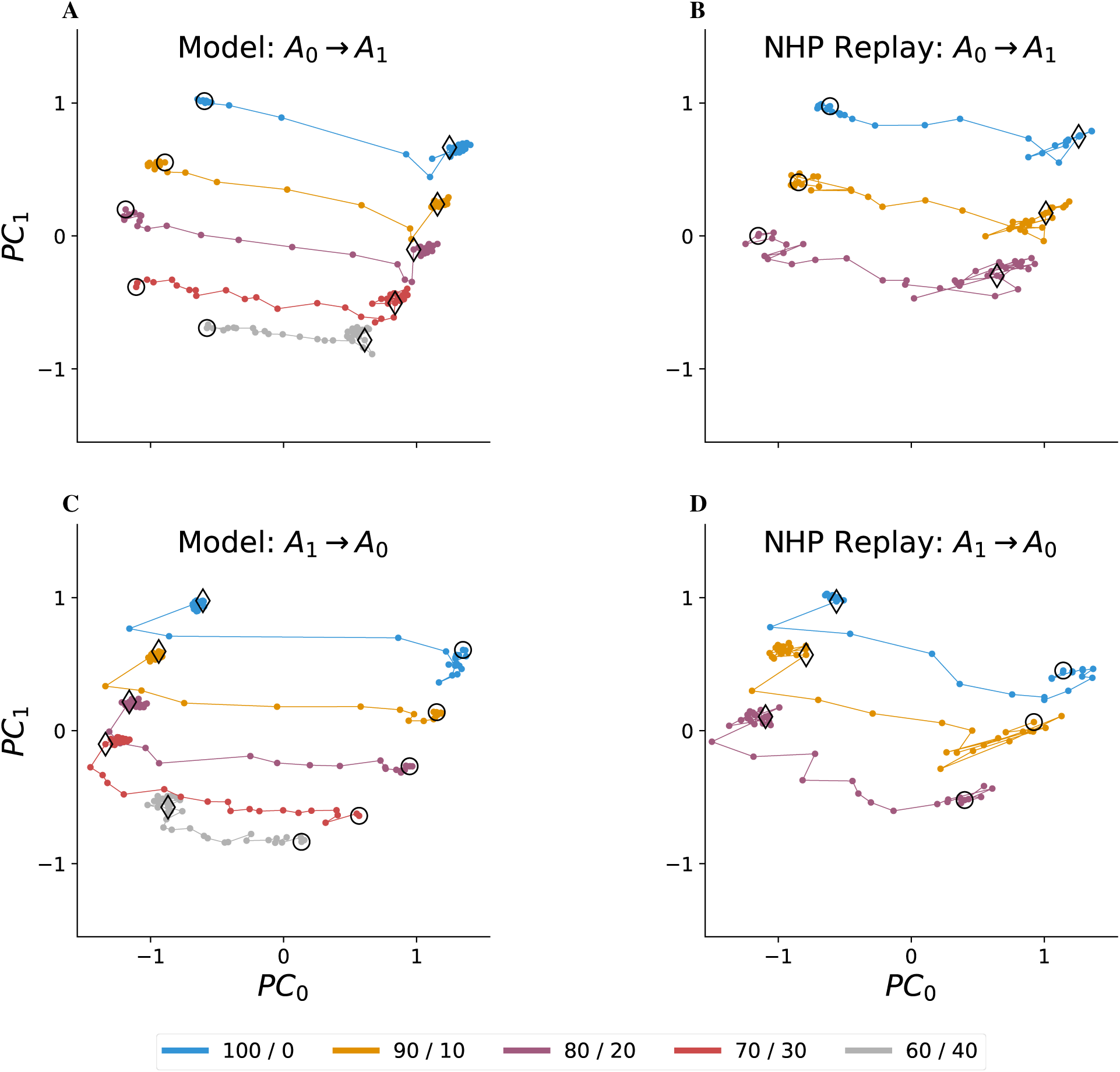
Average belief state for the trials surrounding the time of switch (±19 trials) projected into the first two principal components when the model is determining the actions (A and C) and when the NHP GD actions are replayed into the model (B and D). The trials are split between the transition from *A*_0_ to *A*_1_ (A and B) and the transition from *A*_1_ to *A*_0_ (C and D). Each point represents a different time relative to the switch. The point that corresponds to the time of switch is indicated with a diamond; points that correspond to times before the switch are in the same vicinity as the diamond. The point that corresponds to 19 trials after the switch is indicated by the circle.

These first two principal components capture several key types of information that are necessary to solve this task. First, *PC*_1_ appears to encode the probability scheme, with the deterministic scheme having the highest *PC*_1_ values, and the 60*/*40 scheme having the lowest *PC*_1_ values. Of note is the quick drop in average *PC*_1_ value immediately after the switch that then stabilizes with subsequent trials. This may be due to an ambiguity between a switch occurring and the model evaluating the possibility that it has misestimated the probability scheme. Second, *PC*_0_ encodes the current estimate of highest reward probability action, with *A*_1_ on the left (negative values) and *A*_0_ on the right (positive values). Third, intermediate *PC*_0_ values appear to encode a degree of uncertainty as to which action has the highest probability of reward. In particular, for the deterministic case, the number of trials required to transition from the switch to the vicinity of the circle is relatively small (4 − 5 trials). As the probability of reward decreases, the number of trials required to approach the circle increases (8 − 9 for 90*/*10, and 9 − 10 for 80*/*20). Note that for the 70*/*30 and 60*/*40 schemes, the model is less certain about the action by trial 19; this is consistent with the model not recovering full performance by this trial (see Figure 5A). When the NHP behavior is replayed into the model (Figure 9B,D), the curves show a similar trend. The key difference is the larger variation of the average projected points from one trial to the next. This is due to the smaller number of switches in the averages, and the trial-by-trial variation due to NHP exploration and the stochastic rewards.

### Model Consistency

A key question of any model architecture is the degree of consistency that it displays when faced with different training and evaluation conditions. In addition to the trained model described above, we trained an additional 20 models, each with their own randomly sampled 1024 training and validation sessions. Following training, each model was evaluated with respect to its own validation data set. Figure 10A shows the distribution of percent of correctly chosen actions across the 21 different models, separated by the reward probability. As the reward probability increases, the selection of the correct action increases systematically, reflecting the reduction in ambiguity of the outcome information and the models’ tendency to switch more quickly. In addition, the models show a substantial consistency in their performance for each reward probability, and show significant pairwise differences between neighboring probability levels (*p <* 10^−24^, according to an independent sample t-test).

**Figure 10:**
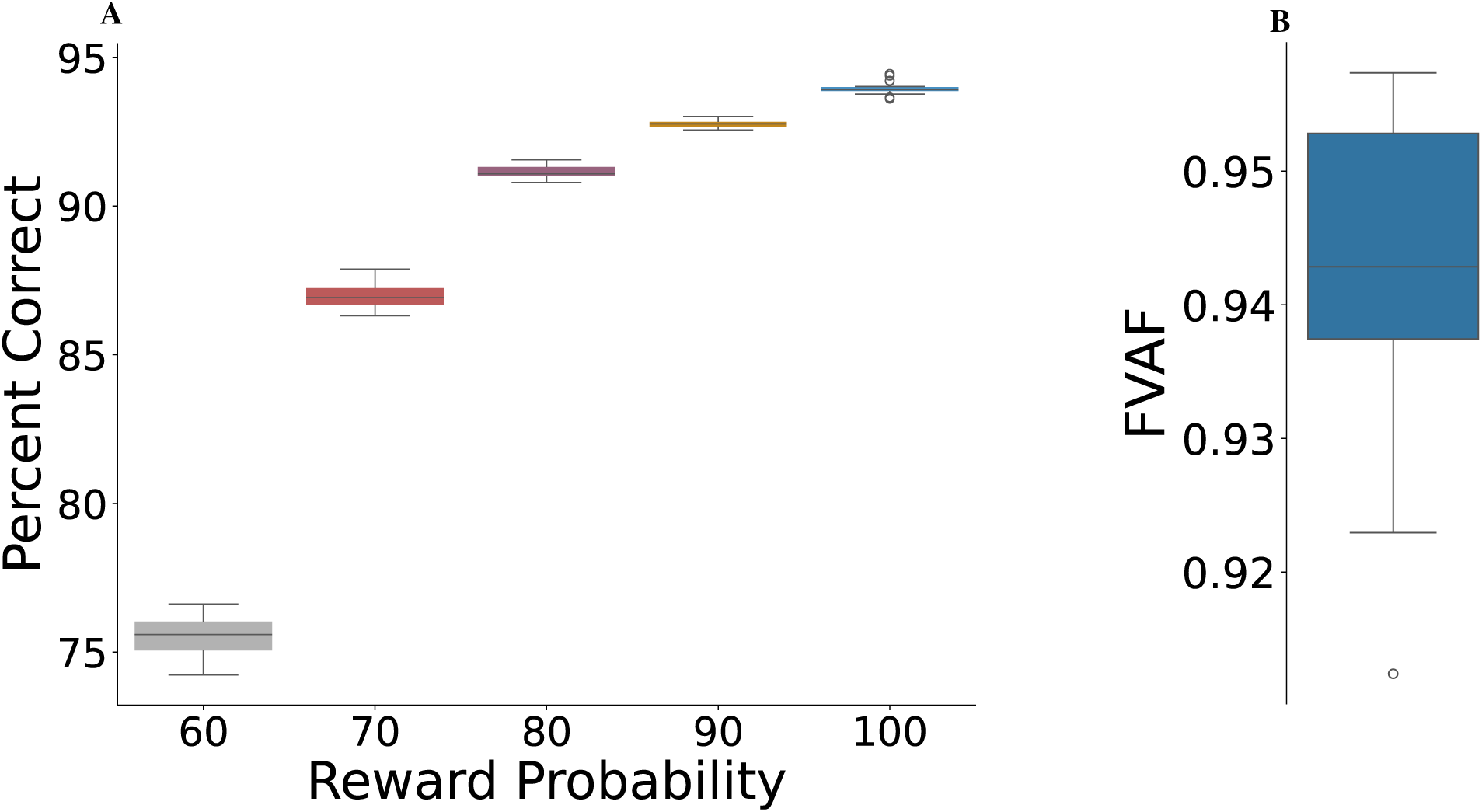
(A) Validation performance as a function of reward probability for 21 independently trained and validated models. Performance increases as the reliability of the reward outcome increases. Each of the models performs very similarly. (B) Fraction of the original model Q-Value variance accounted for (FVAF) by replaying its action/outcome sequences into the remaining 20 models.

Model consistency can also be measured in terms of whether the different models produce similar Q-value estimates, assuming that they experience the same action/outcome sequences. Given the original model, we recorded its estimated Q-values, action choices and the resulting rewards for all of its validation sessions. We then employed experience replay to present these action/outcome sequences to the 20 remaining models, recording their Q-values. Figure 10B shows the degree to which each of the 20 models are able to reconstruct the Q-values estimated by the original model.

### Hyper-parameter Sensitivity

Hyper-parameters are model parameters that are chosen outside of the training process. For the proposed model, one of the key model hyper-parameters is the future reward discount factor, 0 *< γ* ≤ 1. For the reported results, we selected *γ* = 1*/*2, which means that for the purposes of computing *Q*(*X_t_, a*), a subsequent reward of 1 now is scaled to 1*/*2 for the next trial, and to 1*/*4 for the trial after that. Increasing *γ* has the effect of changing the scale of the Q-values, however we found that the behavioral response to the task switch does not change substantially. We interpret this result to mean that the PST task does not require an agent to take into account substantial future horizon to perform well.

## Discussion

Cognitive flexibility is the skill of switching from one task to another, and often requires that the decision to switch be made based on ambiguous information that must be accumulated in time. In this paper, we propose a DRQL-based model of task switching for a probability switching task in which an agent A) must repeatedly select between one of two actions that are probabilistically rewarded, but B) is never explicitly cued as to the reward probability, the correct action to choose at any given time, or the time of task switch. The model consists of modules that A) construct a belief state that integrates the stochastic outcome information across time, and B) evaluates each of the possible actions in terms of the subsequent sequence of expected future rewards (Q-values; see Figure 3).

In contrast to the approaches of Costa et al. (2015) and Lak et al. (2020), belief state update from one trial to the next is implemented using a RNN that is trained in parallel with the action evaluation NN. This approach has the advantage of not requiring a human-designed Bayesian-based belief state update rule, and instead allows the model to learn a representation and update rule that are sufficient to capture just the necessary information to perform well in the overall task. This reduces potential biases that might be implicitly encoded in a hand-crafted design, such as including unnecessary or excluding necessary information from the belief state representation (Qian et al., 2024). Furthermore, for the proposed DRQL-based model, altering the conditions of the overall task (e.g., adding more than two possible actions, or changing the task switching or reward rules) does not require an explicit redesign of the model, only a retraining. This allows for the creation of testable predictions for new tasks prior to use with NHPs (Huang and Rao, 2013).

Because both the RNN and action evaluation components of the proposed model are trained simultaneously, there are no formal guarantees that the DRQL training process will result in an “optimal” solution for belief state and Q-value representation. Nonetheless, we have demonstrated that the model learns to perform the PST consistently across multiple training runs and under different training experiences. In addition, these models learn to produce consistent Q-values, despite these training differences. Together, this evidence suggests that the DRQL approach converges to a common set of solutions for this problem. While the model RNN does not explicitly capture a Bayesian-style representation and conditional update rule, NNs in general have been shown to approximate such processes (Bishop, 1995; Knill and Pouget, 2004; Rao, 2010; Huang and Rao, 2013). In addition, Lambrechts et al. (2022b) present a DRQL-based agent with tasks in which the true full state is known, but that the agent is only given incomplete feedback. They show that the learned belief state captures the true state to a high degree of accuracy.

Behaviorally, the over-trained DRQL model implements task switching through outcome-driven update to the belief state, and does not rely on synaptic changes. A key implication is that the time required to commit to a task switch is not determined by the dynamics of synaptic change, but instead on the dynamics of accumulating uncertain outcome information over time. Specifically, as there is more uncertainty in the reward feedback for each trial, on average, the model requires more trials before committing to the switch. This same time delay effect is seen in the behavior exhibited by NHPs (Figures 4 and 5). These similarities are suggestive of the model agent and the NHP having similar goals during play of the PST (i.e., collecting as much reward as possible).

The ten model neurons that encode the belief state capture different aspects of the agent’s state of knowledge prior to and following the task switch (see Figures 7A-D, S2, S3, and S4). Some neurons explicitly capture the expected probability of reward, while others capture an estimate of which action is the appropriate one to select, and yet others represent different blends of this information. In addition, many neurons respond to the switch at different rates, with the rate often depending on the expected reward probability. Nonetheless, the belief state representation contains redundancies. In particular, most of the belief state variance is encoded within the first few principal components of the ten dimensional belief state. Besides favored action and expected probability, the first two principal components also capture the degree of certainty that the switch has occurred (Figure 9).

Replay of the NHP experience into the trained model provides an approach for asking how the model’s belief state, Q-value, and TD error estimates would evolve under the specific sequence of action choices and rewards experienced by the NHP. We posit that these same types of information will also be encoded in NHP PFC and related neural networks, though the form of the information representation may differ.

TD error is the instantaneous difference between the expectation of the value/quality of selecting an action and the actual outcome that results from that action; here, *actual outcome* encodes a combination of received reward and the expected outcome of future actions (Sutton and Barto, 2018; Watkins and Dayan, 1992). TD error has long been considered as a correlate for DA neural activity (Schultz et al., 1995; Houk et al., 1995; Barto, 1995; Schultz et al., 1997; Qian et al., 2024), though there are acknowledged deviations. Fiorillo et al. (2003) demonstrate a relationship between DA neuron activity and the expected probability of a reward. Nakahara et al. (2004) show that TD error can take time-dependent contextual information into account in learning to interpret cues for potential coming rewards. In our over-trained model, TD error captures the degree of surprise in the received rewards. Under the 100*/*0 probability scheme, TD error is close to zero during task blocks in which the preferred action does not change – even when the model selects the non-preferred action (Figure 4C). When rewards are stochastic, TD error deviates from zero because it is the average reward that is anticipated (panel D). Nonetheless, the average TD error during a block is approximately zero for all but the 60*/*40 probability scheme (Figure 8A; *t* ≤ 0), indicating that the model is largely anticipating the reward probability. It is only the trials following the switch that outcome deviates consistently from expectation. However, this deviation is less for lower reward probabilities because the model had already factored in the expectation of fewer rewards.

## Limitations and Future Work

For the experiments involving replay of NHP-derived experience, we considered only two possible responses by the NHP. In practice, the NHPs do not produce an appropriate response for all trials; in these cases, no reward is given and the NHP is cued to prepare for the next trial. During replay, we removed these non-response trials, implicitly assuming that there is no change to the belief state. However, a non-response action could be explicitly injected by the Task control module (Figure 3).

Action exploration in POMDPs can serve two distinct roles. First, during the learning process, it is important to repeatedly explore the set of action choices across a representative sample of the full belief state. This helps to ensure that both an appropriate belief state representation and the corresponding action evaluation have been discovered. Second, following learning, the choice of an action can serve two roles: 1) gain immediate or future reward, and 2) improve the certainty in the true state so that subsequent actions have a higher likelihood of being rewarded. The *ɛ*-greedy exploration used in the current model, in some sense, conflates these two ideas. A possible resolution to this issue is to make use of an actor-critic style model that replaces the greedy rule with one that first computes a probability distribution across actions as a function of belief state, and then samples actions from this distribution (Konda and Tsitsiklis, 2003; Sutton and Barto, 2018). This would give the model an explicit means of representing the potential need for exploration in order to gain more information, and may better reproduce the NHP tendency to explore more when the probability of reward is low. In addition, rather than minimizing squared TD error, such an architecture could be trained by maximizing the likelihood of the behavior produced by a NHP. This approach would allow for a formal comparison of the models that result from the different objective functions, and could also allow us to account for individual differences in behavioral strategies.

The actions in the proposed model are relatively abstract, capturing concepts such as *saccade to circle* or *saccade to square*. However, there is no explicit representation in the model of *percept*, and the actions are never grounded out to a motor system in terms of specific saccade parameters. We are planning a future model that introduces both, as well as a hierarchy of action decision processes, with the intent of exploring the temporal interaction of subcortical and PFC processes within a DRQL framework.

## Acknowledgements

None

## Author Contributions

- Fagg: Designed research; Performed research; Analyzed data; Wrote the paper.
- Diges: Performed research; Contributed unpublished analytic tools; Analyzed data.
- Rajala: Designed research; Performed research; Analyzed data.
- Habibi: Analyzed data; Edited the paper.
- Suminski: Designed research; Analyzed data; Edited the paper.
- Populin: Designed research; Performed research; Analyzed data; Wrote the paper.

## Supplementary Material

### Individual Monkey Performance

Figure S1A shows the percent of correct response by each NHP around the time of the task switch. NHP GD performed in three different probability schemes: 100*/*0, 90*/*10, and 80*/*20; the number of switches in each curve is 44, 31, and 49, respectively. NHPs M and G only performed in the 80*/*20 probability scheme; the number of switches in each curve is 5 and 25, respectively (small relative to GD). Figure S1B shows the average reward around the time of task switch for each NHP.

**Figure S1:**
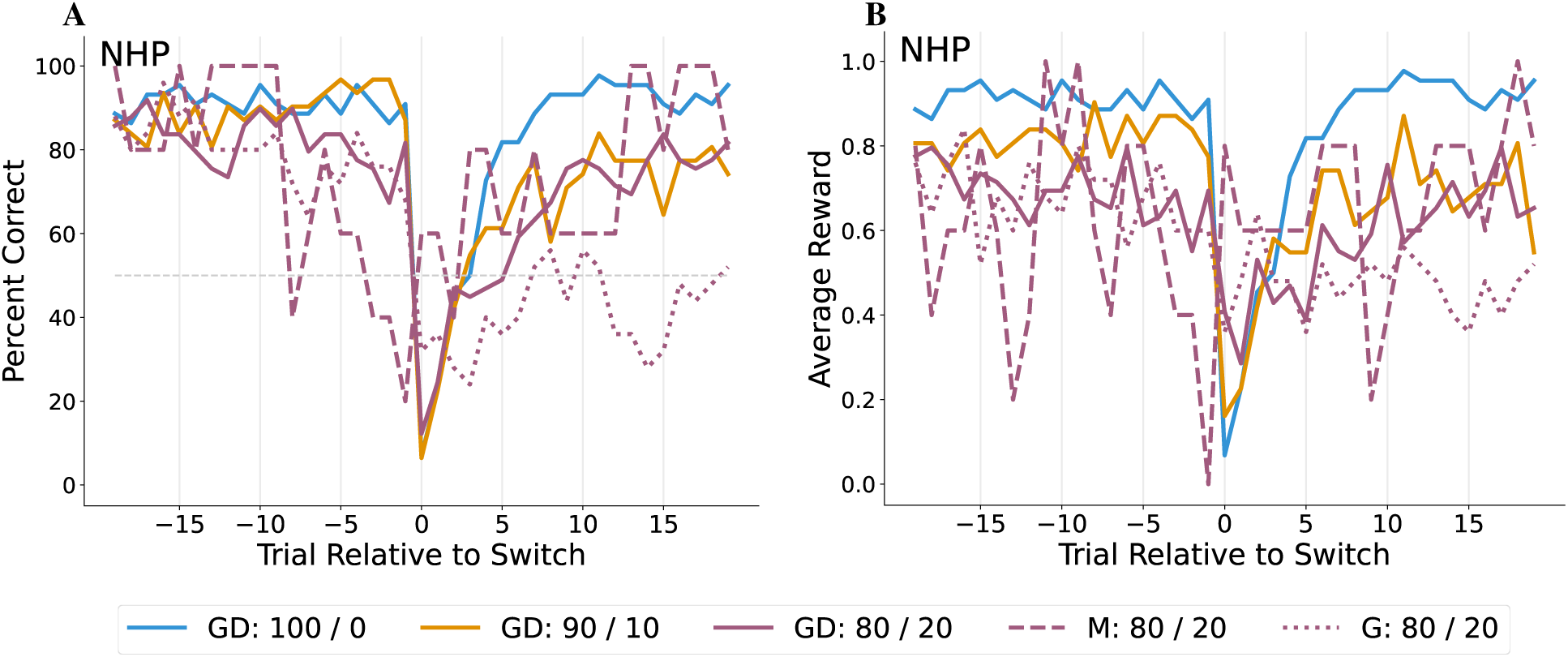
Percent of correct responses relative to the time of switch from one task to the other for NHPs GD, M, and G (A); the average reward relative to the time of switch for the NHPs (B). Trial 0 is the first trial after the switch.

### Belief State

Figures S2, S3, and S4 show the average recurrent neuron activation relative to the time of switch for the model and for when the NHP experience is replayed into the model. These neurons encode different blends of information for the PST, including: best estimate of the correct action to execute, and the probability of reward. For each model neuron, the timing of activation change after the switch varies depending on the probability scheme. Neurons 2, 4, and 5 largely provide an encoding the probability of reward. Neurons 3, 6, 7, and 8 primarily encode the favored action, but are also modulated by the probability of reward. Of these neurons, 6 and 8 exhibit an asymmetry in how probability is encoded; neuron 8 is the extreme, with no modulation by probability for action *A*_1_ leading up to the switch, and no modulation for action *A*_0_ following the switch.

**Figure S2:**
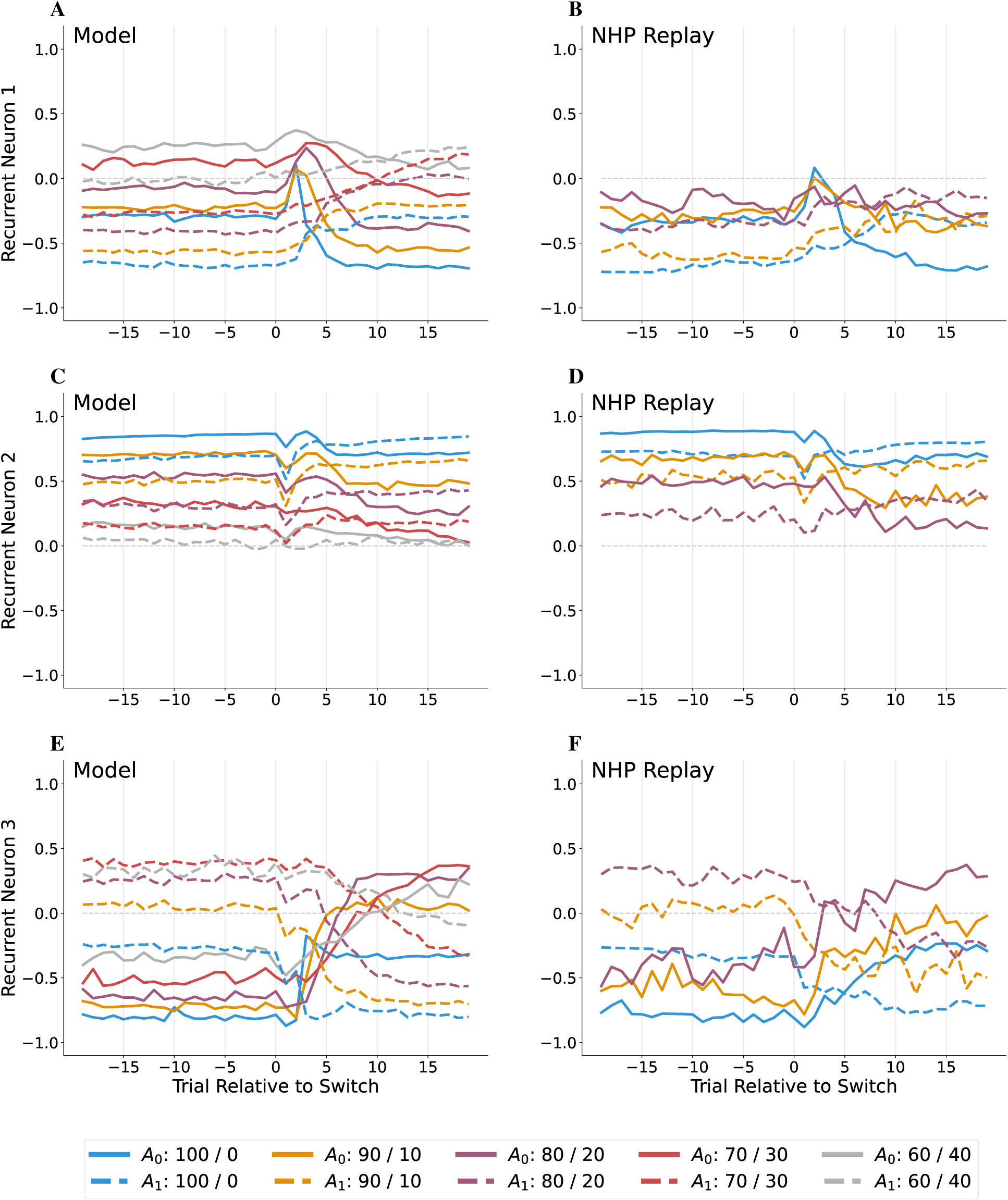
Average model belief state variable response relative to the time of switch for different reward conditions (color) and starting action (*A*_0_=solid; *A*_1_=dashed). Model neurons 1, 2, and 3 when the model determines the actions (A, C, E) and when the NHP behavior is replayed into the model (B, D, F).

**Figure S3:**
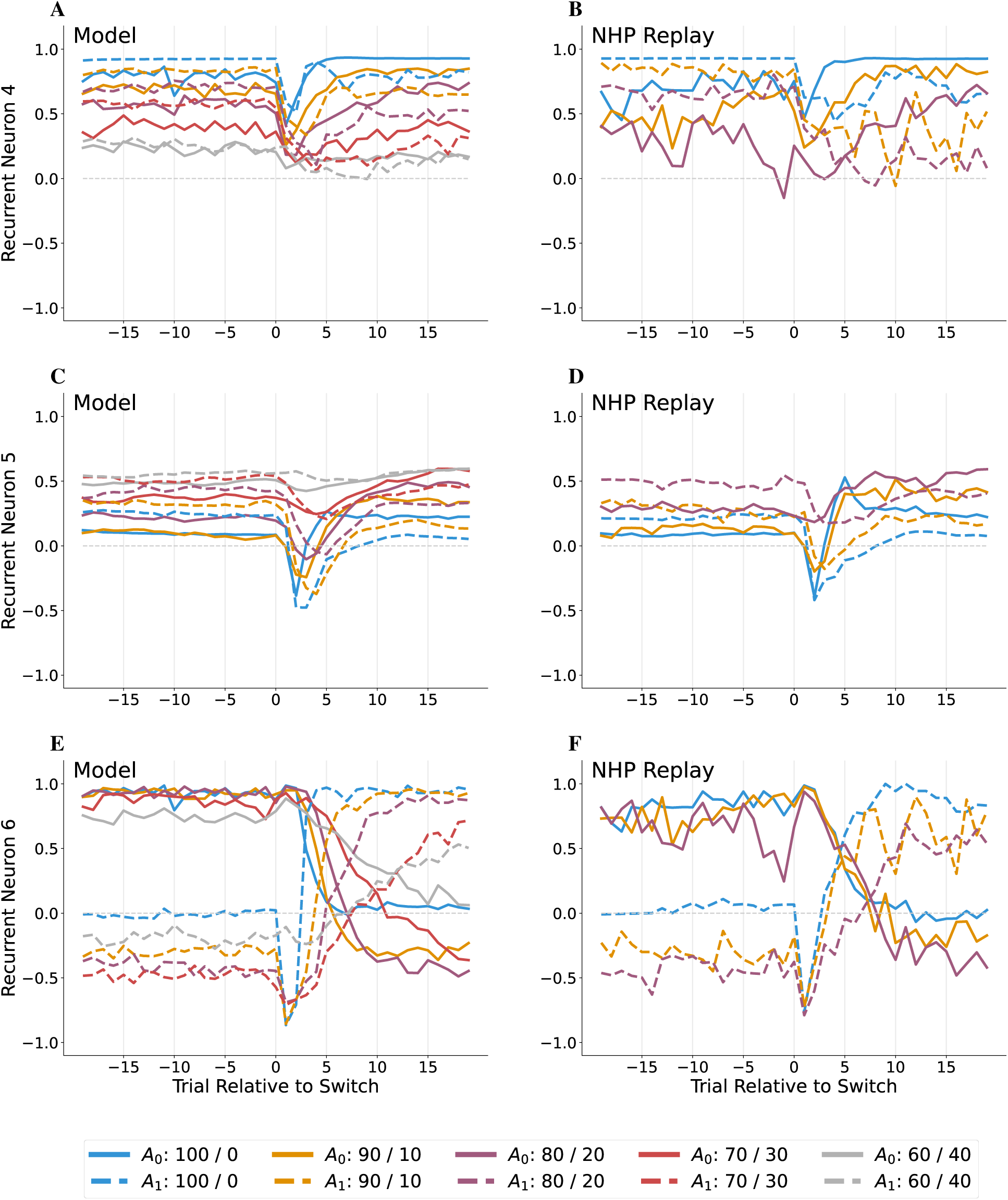
Average model belief state variable response relative to the time of switch for different reward conditions (color) and starting action (*A*_0_=solid; *A*_1_=dashed). Model neurons 4, 5, and 6 when the model determines the actions (A, C, E) and when the NHP behavior is replayed into the model (B, D, F).

**Figure S4:**
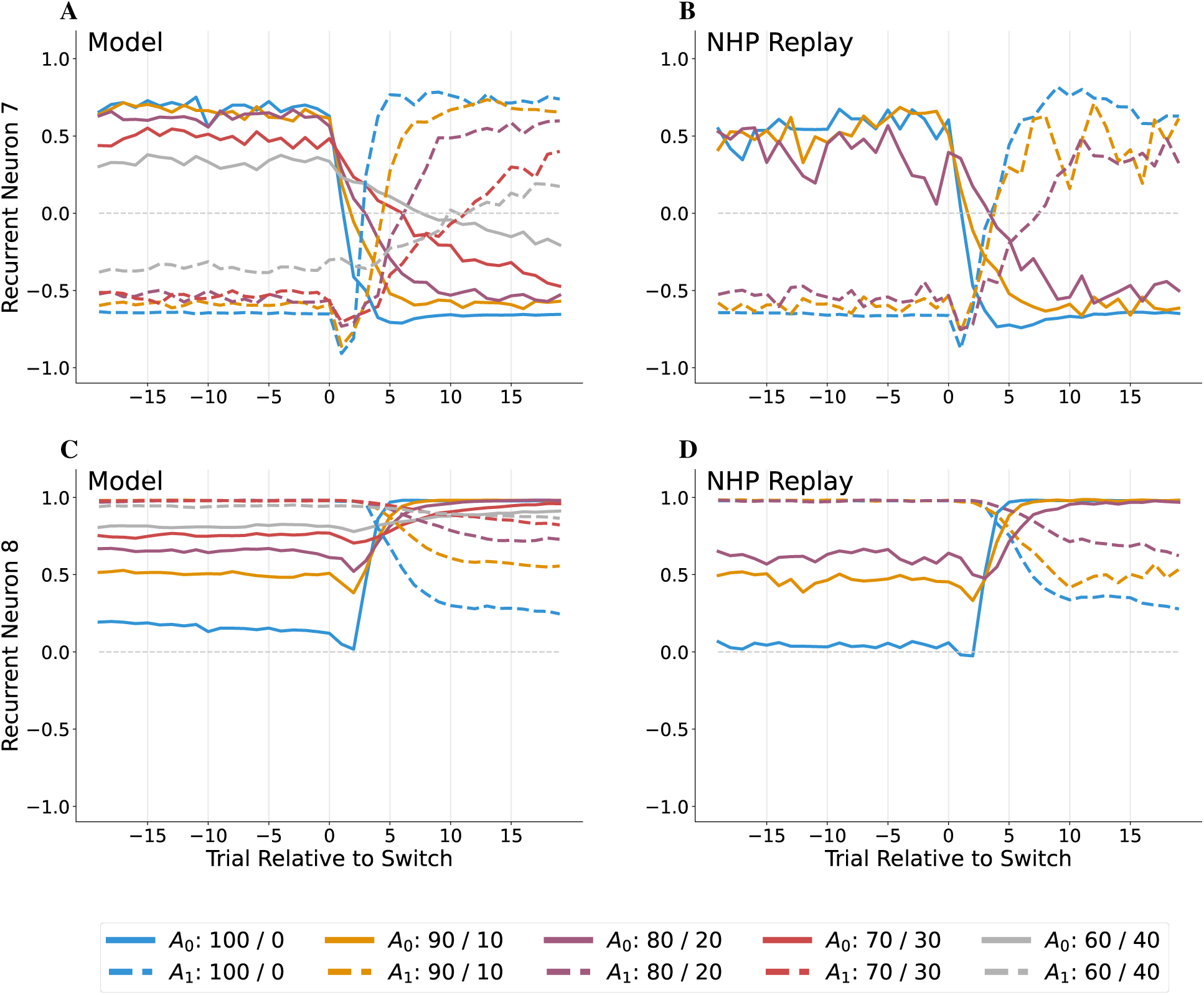
Average model belief state variable response relative to the time of switch for different reward conditions (color) and starting action (*A*_0_=solid; *A*_1_=dashed). Model neurons 7 and 8 when the model determines the actions (A, C) and when the NHP behavior is replayed into the model (B, D).

